# Identification of novel genes involved in phosphate accumulation in *Lotus japonicus* through Genome Wide Association mapping of root system architecture and anion content

**DOI:** 10.1101/600726

**Authors:** Marco Giovannetti, Christian Göschl, Stig U. Andersen, Stanislav Kopriva, Wolfgang Busch

## Abstract

Phosphate is a key nutrient for plants and as it is needed in high quantities. It is highly immobile in the soil and represents a major limiting factor for plant productivity. Plants have evolved different solutions to forage the soil for phosphate and to adapt to phosphate limitation ranging from a profound tuning of their root system architecture and metabolic profile to the evolution of widespread mutualistic interactions, such as those with arbuscular mycorrhizal fungi (AM symbiosis). Despite the prevalence of AM symbiosis throughout land plants, most studies aimed at identifying genes that regulate plant responses to phosphate have been conducted in species incapable of AM symbiosis, such as Arabidopsis. Here we elucidated plant responses and their genetic basis to different phosphate levels in a plant species that is widely used as a model for AM symbiosis: *Lotus japonicus*. Rather than focusing on a single model strain, we measured root growth and anion content in response to different levels of phosphate in a large panel of Lotus japonicus natural accessions. This allowed us not only to uncover common as well as divergent responses within this species, but also enabled Genome Wide Association Studies by which we identified new genes regulating phosphate homeostasis in Lotus. Under low phosphate conditions, we uncovered a correlation between plant biomass and the decrease of plant phosphate concentration in plant tissues, suggesting a dilution effect. Altogether our data of the genetic and phenotypic variation within a species capable of AM complements studies that have been conducted in Arabidopsis, and advances our understanding of the continuum of genotype by phosphate level interaction that exists throughout dicot plants.

**Author Summary:** Phosphate represents a major limiting factor for plant productivity. Plants have evolved different solutions to adapt to phosphate limitation ranging from a profound tuning of their root system architecture and metabolic profile to the evolution of widespread mutualistic interactions, such as arbuscular mycorrhizal symbiosis. Here we elucidated plant responses and their genetic basis to different phosphate levels in model legume plant species, *Lotus japonicus*, a plant commonly used for studying arbuscular mycorhizal symbiosis. We investigated Lotus responses to phosphate levels by combining high throughput root system architecture phenotyping and nutrient measurements with a natural variation approach. We investigated relations between root phenotypic responses and nutrient accumulation and we uncovered, under low phosphate conditions, a correlation between plant biomass and the decrease of plant phosphate concentration in plant tissues, suggesting a dilution effect. By means of Genome Wide Association mapping and integration of multiple traits, we identified new genes regulating phosphate homeostasis in Lotus.

## Introduction

Phosphate is an essential element for plant growth and its bioavailability represents a major limiting factor for plant productivity. Plants coping with phosphate deficiency exhibit dramatic changes at the developmental, nutritional and metabolic levels. For example, in *Arabidopsis thaliana,* the root developmental program has been described to be highly affected by phosphate deficiency. Primary root growth of the reference accession Col-0 is inhibited and there is an increase of lateral root formation and root hair growth (López-Bucio et al., 2002). The key genetic determinants of this process have been identified, mainly through forward genetic screening (Svistoonoff et al., 2007; Wang et al., 2019). Recently, it has been shown that a main driver of primary root growth arrest is toxicity of iron that, upon phosphate starvation, accumulates in the meristematic zone and induces a progressive loss in the proliferative capacity of the cells, causing reduction in meristem length (Müller et al., 2015). Plants facing phosphate starvation exhibit a dramatic remodeling of main cellular processes: a transient DNA methylation altering gene transcription (Secco et al., 2015), mainly driven by the phosphate-dependent interaction of SPX1/PHR1 (Puga et al., 2014). In addition, phosphate-depleted plants usually display a high turnover of phospholipids into sulfolipids (Essigmann et al., 1998). Furthermore, there is a substantial interaction of phosphate related processes and other environmental factors. For instance, red light frequencies lead to an increase of phosphate uptake and Arabidopsis accessions with light-sensing defects, such as Lm-2 and CSHL-5, take up less phosphate (Sakuraba et al., 2018). Moreover, phosphate accumulation is altered by the abundance of metal ions, such as iron and zinc, with the extent of the impact being dependent on the genetic background (Briat et al., 2015; Kisko et al., 2018). Taken together, responses to phosphate and phosphate homeostasis in plants are regulated in a complex manner and are substantially dependent on plant genetic diversity and environmental abiotic and biotic factors.

For biotic factors, it was recently shown in Arabidopsis and related plants how fungi and bacteria play a key role in mediating phosphate accumulation (Almario et al., 2017; Hiruma et al., 2016). Molecular hubs of phosphate metabolism, such as PHR1, can define the composition of microbial communities (Castrillo et al., 2017) and the analysis of microbial contribution to plant phosphate accumulation can even be mathematically modeled, giving rise to the possibility to design ad-hoc microbial communities with desired effects on plant phosphate uptake (Paredes et al., 2018). The most prominent example of microorganisms providing plants with phosphate is the symbiosis of plants with arbuscular mycorrhizal (AM) fungi. A key aspect of this is that fungal hyphae are very efficient in exploring the soil and taking up the highly immobile phosphate. Consequently, up to 80% of the needed phosphate can be acquired and transferred to plants by these fungi (for up-to-date reviews Choi et al., 2018; Lanfranco et al., 2018; MacLean et al., 2017). Numerous plant genetic determinants that are needed for the establishment of a functional mycorrhizal symbiosis have been described over the last 20 years but a lack of high-throughput methods and the complexity of the system impaired the comprehensive understanding of the crucial and complex feedback happening between plant phosphate status and the establishment of AM symbiosis (Carbonnel and Gutjahr, 2014).

Most genetic screens aiming for identifying genes that regulate plant responses to phosphate have been conducted in species incapable of AM symbiosis such as Arabidopsis. In this study we targeted the genetic basis of responses to phosphate levels in a plant widely used as a model for AM symbiosis. To do so, we conducted large-scale studies of *Lotus japonicus* natural variation of root responses to different levels of phosphate, coupled with the measurements of anion accumulation. We discovered profound correlations of plant size and plant phosphate concentration that should be taken into account when working with concentration measures. Using high-density SNP data from the 130 Lotus accessions, we conducted Genome Wide Association Studies (GWAS) for all measured traits, finding hundreds of genetic loci associated with variation in phosphate-related traits. Finally, by comparing the lists of candidate genes for root system architecture and phosphate accumulation, we identified a Leucine-Rich Receptor kinase and a cytochrome B5 reductase involved in phosphate homeostasis as high confidence causal genes, which was further corroborated by phosphate dependent phenotypes of loss of function mutants for these genes.

## RESULTS

### Phosphate deficiency shapes natural variation of root growth and anion levels in roots and shoots of *Lotus japonicus*

To study the genetic bases of root responses to low phosphate and phosphate accumulation in *Lotus japonicus* (Lotus) tissues, we performed a detailed root phenotyping of a panel of 130 diverse Lotus natural accessions (Shah et al., 2018) over a 9-day time course and subsequently quantified the anions of the main macronutrients from the same material (Fig. 1a). In particular, we grew the plants on vertical plates for 9 days on a modified Long-Ashton medium (Supplemental Table 1), containing either 20 μM (LP) or 750 μM (HP) of phosphate. We scanned the plates daily, at the same time of the day and at the end of the 9th day, we harvested and weighed total root and total shoot (stem and leaves) material for subsequent quantification of main macronutrients: nitrate, phosphate and sulfate.

**Fig. 1.**
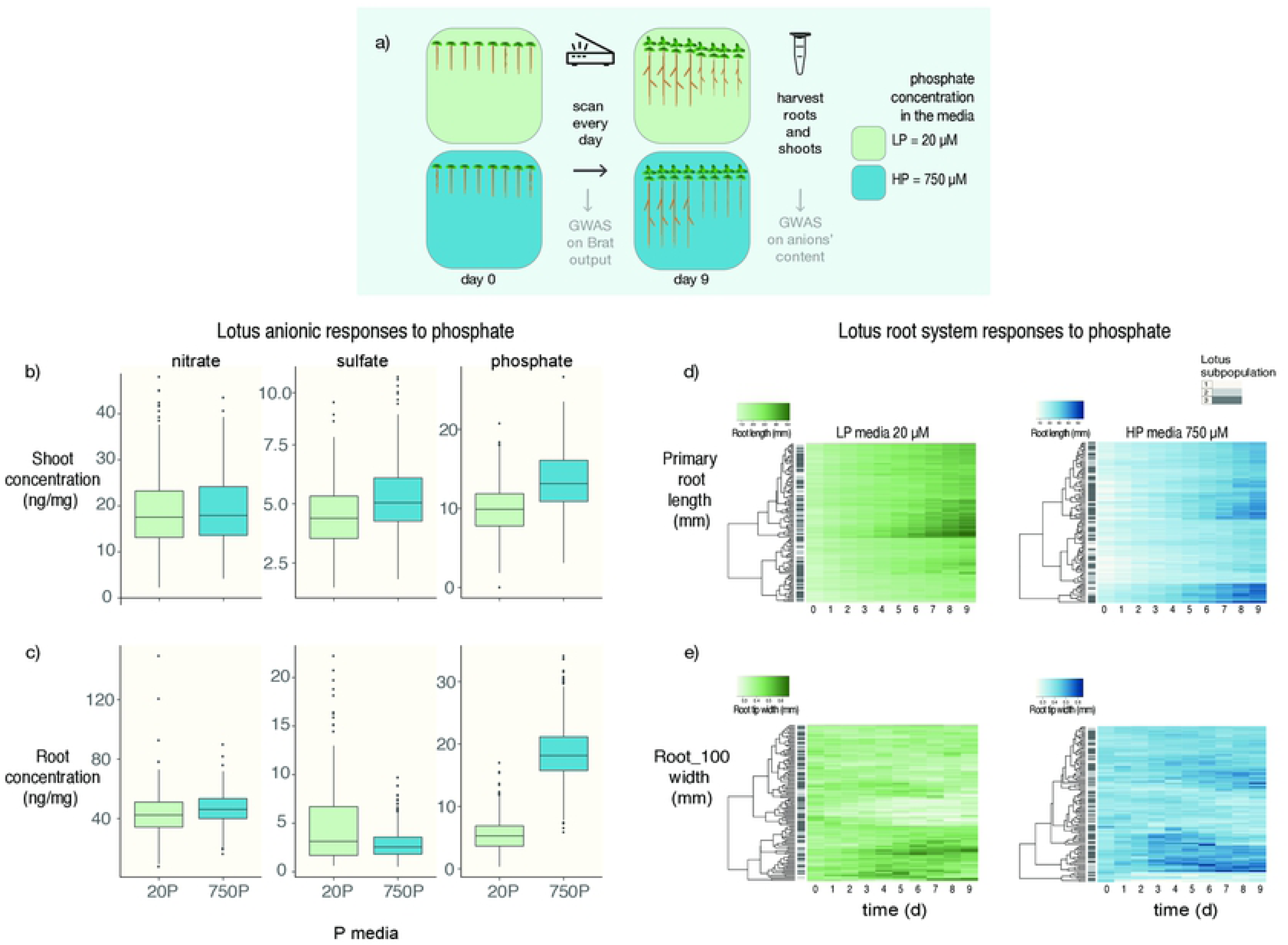
Lotus japonicus natural variation of root responses to phosphate media levels over time. a) Set up of the experiment. One-hundred and thirty Lotus accessions were grown on low (20 µM) or high (750 µM) phosphate media for 9 days. Plates were scanned every day and root traits were quantified and segmented using Brat. After 9 days, shoots and roots were harvested and nitrate, phosphate and sulfate concentration were measured. All the traits were then used for running GWAS. Here, we show representative traits of anions’ accumulation and primary root traits of the Lotus panel. b-c) Concentration of anions in roots and shoots depends on the phosphate concentration on in the media. Both shoot and root accumulation of anions’ shows a significant effect of phosphate concentration in the media. A strong effect is shown by phosphate content in roots and shoots and sulfate content in shoots. d-e) Over a 9-day time course, Lotus natural access ons show a high diversity in root growth over LP and HP both for primary root length and root width. Clustering of responses to nutrient is not dependent on Lotus subpopulation origin.

As shown in Figure 1b-c, wide variation among macronutrient concentrations is observed in different accessions. Phosphate levels in the medium not only affect plant phosphate concentration in roots and shoots, but also plant sulfate levels and, to a minor extent, nitrate concentration in roots, exposing a similar cross-talk between the three anions in Lotus as the one that had been described in Arabidopsis (Kellermeier et al., 2014). With the exception of phosphate concentration of plants grown under phosphate starvation, the concentration of these anions did neither depend on plant size nor on plant developmental stage (Supplemental figure 1).

By using a modified version of the Brat Fiji plugin (Giovannetti et al., 2017; Slovak et al., 2014), we quantified 16 root traits per each day. Root traits showed a broad spectrum of responses among accessions (Fig. 1d,e). There was a pronounced effect of phosphate on the majority of traits and interactions of effects between genotype and phosphate (Supplemental file 1). We then explored whether the response to phosphate levels merely reflects the genetic relation between Lotus natural accessions. For this we conducted hierarchical clustering of root growth and root tip width in both phosphate levels and found that the clusters do not reflect the established genetic Lotus subpopulation structure (Shah et al., 2018) but rather depend on phosphate level in the medium (Fig. 1d,e). In contrast to the early root growth responses described in the Arabidopsis reference accession, low phosphate medium does not induce a dramatic arrest of primary root growth in most of the Lotus accessions (Supplementary Fig. 2a). Similar PR responses have also been also observed in non-reference Arabidopsis accessions (Chevalier et al., 2003), the diversity of root growth responses to phosphate levels in Lotus seems to resemble that in Arabidopsis. Beyond total root length, root traits related to the root diameter, such as root width, are substantially influenced by phosphate deficiency: the width of the first 20% adjacent to the root/shoot junction (denominated as trait “Root width 20”) and the distal 20% (“Root width 100”), do get larger over time in plants grown in HP (Suppl. Fig. 2 b,c).

**Fig. 2.**
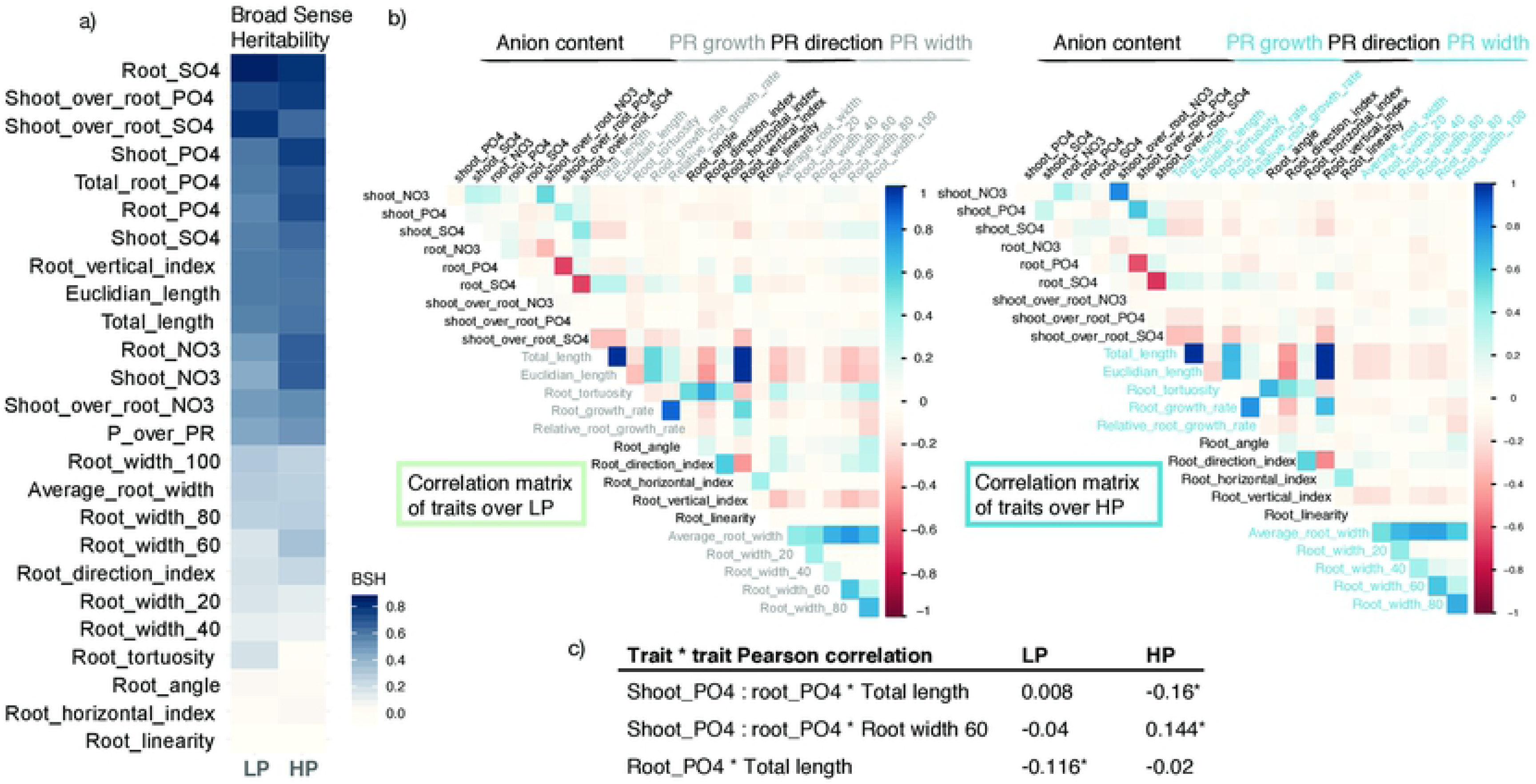
Pattern of Lotus correlation among root and anions’ traits and broad sense heritability at day 9 under LP or HP. a) Broad Sense Heritability of all measured traits on low and high phosphate. Highest heritability is shown by traits related to nutrient accumulation. Among root system architecture traits, those related to primary root length show higher broad sense heritability b) Heatmap of Pearson’s pairwise correlations on every measured trait (root system architecture and nutrient accumulation) on day 9. On a population level, the traits acquired by Brat show distinct and recursive features: root width from the different root parts are all positively correlated. A similar pattern is shown also by traits related to primary root growth, such as total length, Euclidian length, root vertical index. By contrast root width and length parameters are negatively correlated: the longer the root, the finer it gets. Correlation between root system architecture traits and tissue anions concentration show the same pattern in LP and HP with a few exceptions. c) Total length and shootP04:rootP04 (phosphate translocation) do show an opposite behavior or among the two conditions. In particular in low phosphate plates, primary root length is negatively correlated with root phosphate concentration in discordance with normal phosphate condition. By contrast, shootP04:rootP04 has a clear opposite pattern. Pearson’s r value are indicated and the asterisk represents p value 0.05

The broad sense heritability (BSH) was different among the traits that we measured. While the variation of some traits could not be explained by genetics (Root linearity and root angle, ∼0%), most traits are genetically determined and some to an extraordinarily high degree (root_SO4, 86%). Generally, traits related to nutrient accumulation showed higher heritability (Fig. 2a). Among the root developmental traits, total root length showed the highest BSH (∼60%), consistently with data from other root system architecture studies in *Arabidopsis thaliana* (Ristova et al., 2018).

Taken together, we found that Lotus natural accessions exhibit a great variation of responses to phosphate concentrations, both at the phenotypic and at metabolic level. Moreover, the similar profound extent of natural variation of root growth between Arabidopsis and Lotus suggests that a large genotype by phosphate level interaction exist within species throughout the dicot group and regardless of whether a species is capable to form AM symbiosis.

### There is a trade-off of root phosphate concentration and root length specifically under low phosphate conditions

Phosphate starvation has a profound impact on root development and nutrient accumulation. Nevertheless, the link of the two processes remains largely unknown. Since we quantified phosphate, sulfate and nitrate concentrations from roots and shoots of single plants, and also measured root traits over time, we were able to compare trait correlations in LP or HP. This represents a valuable dataset to investigate the links between plant developmental adaptations and cellular metabolic tuning. For this, we calculated the pairwise Pearson’s correlation coefficients of all root trait data and anion content in high and low phosphate from all Lotus accessions (Fig. 2b). The contrasting phosphate levels in the two media did not perturb the majority of correlations among traits (Fig. 2b): for example root width at different sectors along the root were highly positively correlated regardless of phosphate level, as well as the traits related to root length (Euclidian length, root growth rate, total length and relative root growth rate). Root length was negatively correlated with root width, in both conditions, indicating that longer roots are usually thinner in our working conditions.

Nevertheless, we could observe several peculiar correlations occurring exclusively in one of the two conditions: first, the ratio of phosphate concentration between shoot and root, that describes how much of the uptaken phosphate is transferred to the shoot, is significantly negatively correlated with root length and positively correlated with root width under HP but not LP (Fig. 2c). This indicates that under HP less phosphate is allocated to the shoot when roots are elongating. However, when phosphate becomes limiting in the medium (LP), the total length of roots and root phosphate concentration are moderately negatively correlated (Fig. 2b and supplementary fig. 3). This suggests a model in which under LP the available phosphate is distributed over a larger amount of root tissue if roots are longer and is thereby diluted. This dilution model would furthermore also explain the negative correlation between plant biomass and phosphate concentration (Suppl figure 1). By contrast, and in agreement with this dilution model, the negative correlation of phosphate levels and root length is completely absent in HP media (Fig 2c) and much reduced when considering plant biomass and phosphate concentration (Suppl figure 3).

**Fig. 3.**
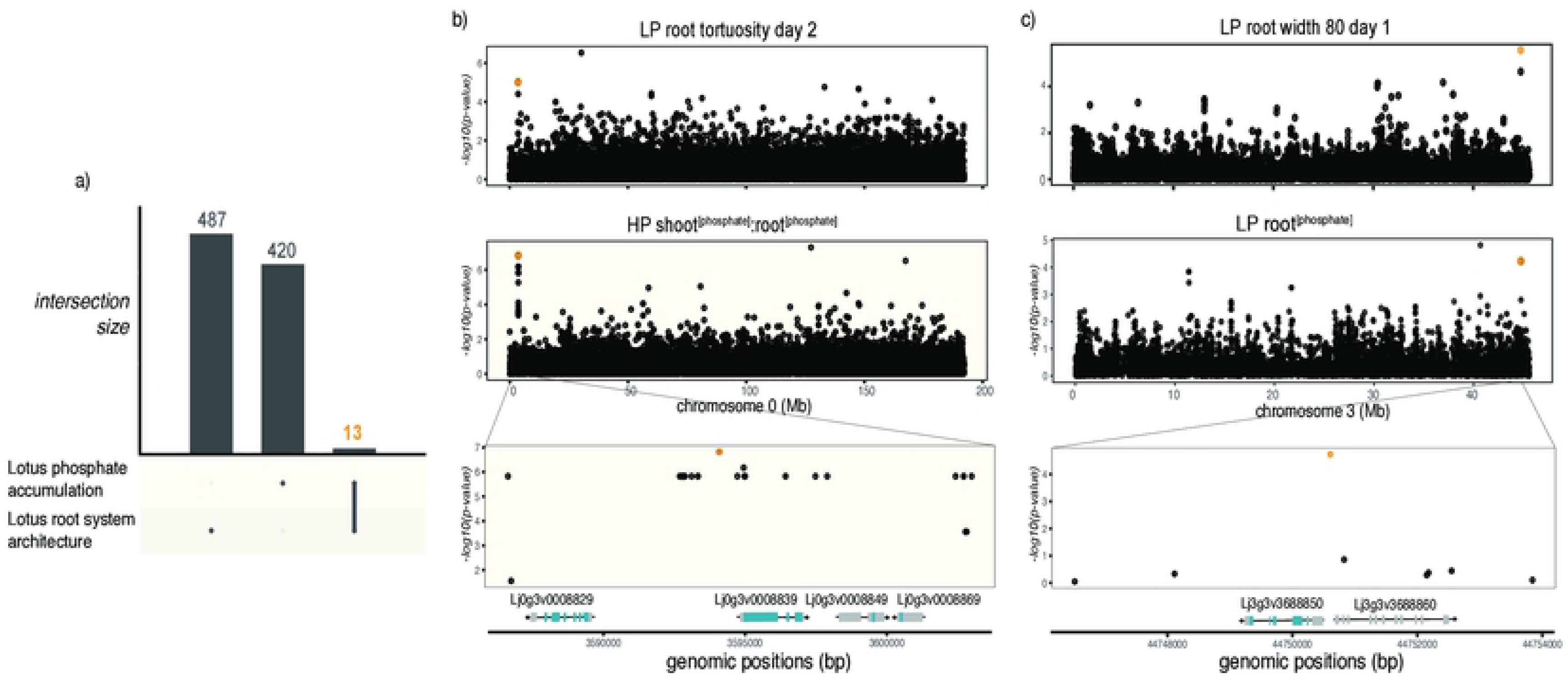
Overlap between GWAS hitsfrom root system architecture and phosphate accumulation. a) The intersection of candidate genes associated with changes in root system architecture (487) and phosphate accumulation traits (420) is 13, corresponding to 7 genomic regions. B) Manhattan plots of two traits leading to the identification of one genomic region on chromosome 0. A close-up on that region shows a 20kb region containing two protein-coding genes and two non-coding genes. c) Manhattan plots of the two traits leading to the identification of a genomic region on chromo some 3. A close up on a 20kb region containing one single protein-coding genes, a cytochrome B5 reductase

Altogether a parallel quantification of phosphate accumulation and root system architecture of a panel of 130 Lotus grown at two different phosphate concentrations allowed for the detection of trait correlation structure, showing that most of trait-correlation are not perturbed by phosphate levels. Our analysis also revealed that, in our experiment, the majority of nutrient accumulation traits show higher broad sense heritability compared to root developmental traits.

### GWAS for Lotus phosphate related traits identifies hundreds of unknown and known candidate genes for phosphate homeostasis

Given the extensive natural variation of most traits in the Lotus panel, we conducted a Genome Wide Association Study (GWAS) for each trait and each time point in LP and HP condition, using a mixed model algorithm, corrected for population structure (Yu et al., 2006; Kang et al., 2008; Seren et al., 2012) and using sequencing-based SNPs for the Lotus accessions (Shah et al., 2018). Each trait led to the identification of genetic loci significantly associated with variation of those traits: using a Benjamini-Hochberg FDR threshold of 5%, we found 900 SNPs associated with root growth parameters of plants grown on LP media (Supplementary table 3), 939 SNPs with HP conditions (Supplementary table 4), 3673 of LP/HP root growth ratio (Supplementary table 2), 104 associated with phosphate content and 220 with anion content (Supplementary table 5 and 13).

Several obvious candidate genes were in these lists. For genes associated with morphological root traits, our analysis identified homologues of several known regulators of root development, such as an homologous gene of SCARECROW (Di Laurenzio et al., 1996), Lj3g3v0821320, associated with Root horizontal index at day 6 under low phosphate conditions. Similarly, sequence variation in the genomic region of a Lotus BIG BROTHER homologue (Cattaneo and Hardtke, 2017), Lj3g3v0489450, is associated with relative root growth rate over day 1-2. Another candidate gene identified as a potential regulator of Lotus root responses to low phosphate is the homologous gene of STOP1 (Lj0g3v0231229) that is associated with Root width 20 variation at day 3. In Arabidopsis, STOP1 was recently described as a key regulator of early root responses to phosphate deficiency-induced iron toxicity (Mora-Macías et al., 2017; Balzergue et al., 2017), therefore a similar role is conceivable for Lotus. Various fatty acyl-CoA reductases are highly associated with root width 80 day 2 and have been shown to be involved in alcohol synthesis as response to various stress and suberin accumulation (Domergue et al., 2010). In parallel, the quantification of phosphate concentration at root and shoot level for the Lotus natural accessions grown at two different levels of phosphate led to the identification of several genetic loci associated with variations in those traits. Among the genes associated with phosphate concentration-related traits, we identified a trehalose-phosphate phosphatase-like protein (Lj4g3v2820240) associated with shoot_[PO4]_:root_[PO4]_ in LP. A significant association was also detected for a SNP within a UDP-glucuronic acid decarboxylase gene (Lj4g3v2312430), associated with shoot phosphate concentration in HP plants. Variation in the genetic region spanning a candidate sugar/phosphate translocator (Lj1g3v4830440) is correlated with root phosphate concentration in HP, as well as a UDP pyrophosphate phosphatase (Lj0g3v0276539) associated with shoot phosphate concentration in HP. Altogether many genes related to phosphate recycling seem to be linked with phosphate accumulation in shoots and roots in both media conditions, even though GO enrichment analysis does not highlight any obvious category (Suppl. Table 11 and 12).

### Overlap among Lotus GWAS from different traits exposes loci associated with both Lotus root growth upon phosphate starvation and phosphate accumulation

Beyond investigating specific genetic associations between Lotus SNPs and particular root traits or phosphate accumulation values, one of our main interests within this study was to use both our metabolic and root growth data to specifically determine genes that control Lotus responses to phosphate. To accomplish that, we considered all genes in 10 kb genomic regions (Linkage Disequilibrium decays in Lotus, r^2^ <0.2, with 10 kb (Shah et al., 2018)) centered on the SNPs passing our GWAS detection threshold. Because it was our purpose to assess overlaps, we took a non-conservative threshold for this approach. We considered up to 500 SNPs with a Benjamini-Hochberg threshold of 10^-5^ for the two groups of traits: phosphate content and root system architecture. This approach led to an overlap consisting of only 7 genomic regions (Supplemental table 8) that were associated with both root and phosphate accumulation traits. These regions were in close proximity to 13 genes (Fig. 3a). We focused on two of these regions, each encompassing a single protein coding gene that was expressed in roots (according to publicly available expression data (Mun et al., 2016) and had possible functions related to signaling and/or acquisition of phosphate. One locus was associated with shoot_[PO4]_:root_[PO4]_ on HP and root tortuosity at day 2 on LP (Fig. 3b and supplementary figures 8 and 9) and the other locus was associated with root phosphate concentration on LP and root width 80 at day 1 on LP (Fig. 3c). Interestingly, this locus was also associated with LP:HP root growth rate between day 7 and day 8 (Suppl. table 2). To further validate an existing interaction among these two pairs of traits, we also performed a multitrait GWAS, based on LIMIX (Casale et al., 2015; Turley et al., 2018), a mixed-model approach enabling analysis across multiple traits while accounting for population structure. Consistently with our overlap analysis, the same loci are associated with both traits for which they were detected (Suppl. Fig 10 and 11), therefore constituting suitable candidate genetic regions associated with Lotus root responses to phosphate.

In both cases, the above mentioned SNPs span a 10kb LD region with exclusively one protein-coding gene that is expressed in roots: Lj0g3v0008839, coding for a LRR receptor-like serine/- rich (LRR-RK) protein on chromosome 0 and Lj3g3v3688850, a putative cytochrome b5 reductase (CYT). To test whether these candidate genes have a role in controlling responses to phosphate level, we selected multiple insertional mutants for each gene from Lotus Base (Małolepszy et al., 2016; Mun et al., 2016) as represented in Fig. 4a. We quantified the tissue phosphate concentration in homozygous mutant plants for both genes in three different phosphate concentrations (20, 100 and 750 μM), 10 days after transfer to specific media plates (Figure 4b,c). All three independent mutant lines of the LRR-RK showed an increased total plant phosphate concentration on high phosphate concentrations (Figure 4b,c and Supplementary table 7). Accordingly, we named the gene *LAMP* (**L**RR-RK **A**ccumulating **M**ore **P**hosphate). Two out of three CYT mutant lines showed an increased total plant phosphate concentration on high phosphate concentrations, specifically driven by shoot phosphate levels (Supplementary figure 5). The WT-like responding mutant line was different in the transposon insertion site with respect to the other two mutant lines and still might have some remaining activity of the protein due to transposon insertion at the far end of the gene (Fig. 4a). While the observable effects of the loss of function of these two genes was significant in HP, a diverse and variable phosphate accumulation took place in LP conditions. We reasoned that the effect of LP on biomass and root length that we had observed earlier (Suppl. Fig. 1) might confound the effects on phosphate content. Therefore we assessed these correlations also across the wt and LORE1 insertion lines. As found within the large panel of accessions, we observed that total phosphate concentration from mutant and wt plants grown under strong or mild phosphate starvation (20, and 100 μM) was strongly negatively correlated with plant biomass (ρ=-0.58, ρ=-0.55, respectively) (Fig. 5), but this correlation is completely absent under HP (750 μM). Interestingly the stronger correlations are more pronounced considering the whole plant compared to root or shoot separately (Suppl. Fig. 6). While we selected these two candidate genes for being associated with phosphate level dependent phosphate content as well as root growth traits, we could not observe a consistent and significant root growth difference to wildtype in any of the tested phosphate concentration (Suppl. Fig 7).

**Fig. 4.**
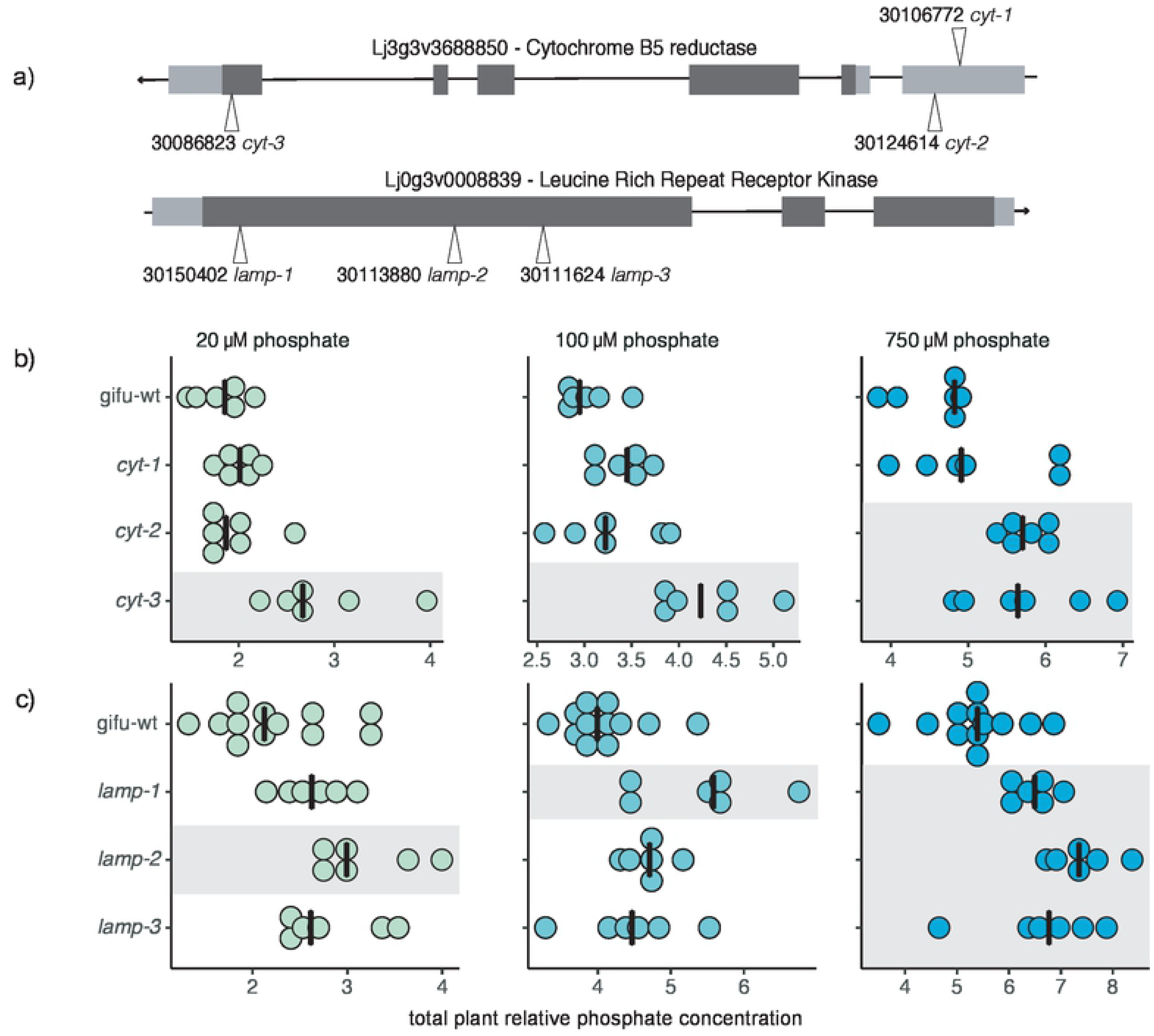
Cytochrome B5 reductase and LRR mutants accumulate more phosphate than wt in high phosphate media. a) Gene structure and insertional mutants used in this study Each number represents the Plant ID from Lotus Base. Three insertional mutants per each gene were used. b) Plant phosphate concentration levels of wt and LORE1 cytochrome B5 reductase insertional mutant plants growing under low {20 µM) or mid (100 µ M) or high phosphate level (750 µM). Whereas at low and mid concentration, only cyt3 show a signficant difference compared to wt plants, at high phosphate concentration also cyt2 is accumulating significant y more phosphate. c) Total phosphate concentration of LRR-RKmutant plants and wt in the three phosphate media conditions. Both three LORE1 insertions show a h gher phosphate accumulation compared to wt. Each dot represents a single plant and black vertical lines represent the median among the group. Levels of phosphate are expressed relative to wt root plants at 20 µM. Different shades represent different groups compared to wt, following Anova test on estimated marginal means (Tukey’s adjusted p value < 0.05).

**Fig. 5.**
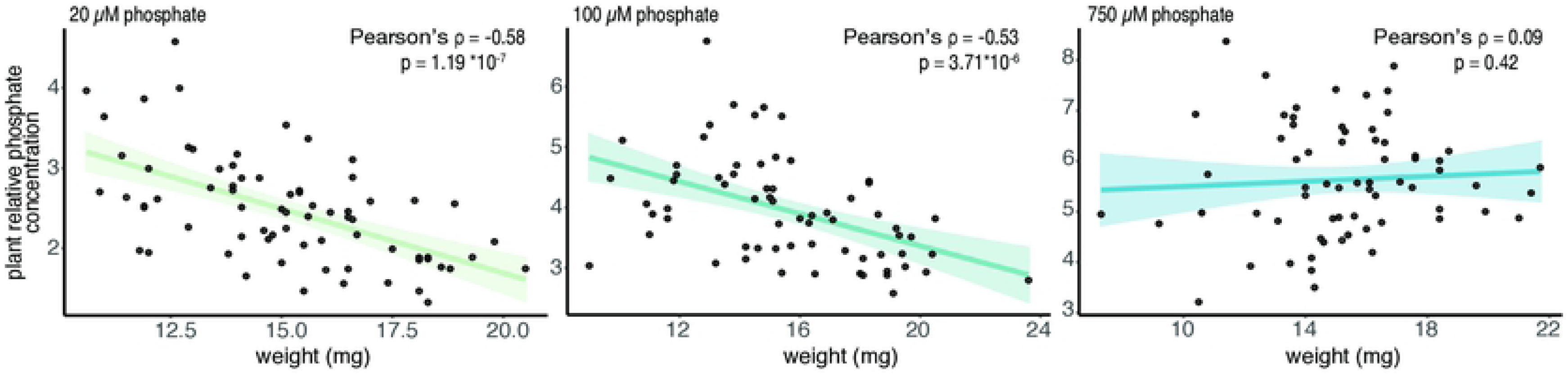
Plant phosphate concentration is negatively correlated with plant biomass when phosphate is the limiting factor. Phosphate concentration levels of plants growing under low (20 µm) or mid (100 µM) phosphate is highly negatively correlated with plant biomass (Pearson’s correlation is −0.58 and −0.53 respectively and p-value <10-6). By contrast under high phosphate level (750 µM), no significant correlation has been observed between plant biomass and plant phosphate concentration. Each dot represent a single plant from different experiments. Phosphate concentration is calculated relative to wt roots phosphate concentration under 20 µM. Colored lined and colored shades represents linear regression and 95% confidence intervals.

Altogether, we provided the community with an atlas of root growth and anion related Lotus phenotypes, showed that root system variation within a genotype by phosphate interaction is not specific to Arabidopsis but also happens in a plant able to form AM symbiosis-even in the absence of the symbiont-, we generated a catalogue dataset of genes associated with root and metabolite responses to phosphate, we investigated phenes and cross-links shaping Lotus natural variations of responses to phosphate and we genetically validated new candidate genes involved in phosphate accumulation. Lastly, a clear confounding element has been unveiled that could prevent future inappropriate conclusions.

## Discussion

In this study, we generated a comprehensive atlas of root system architecture and nutrient accumulation responses to two levels of phosphate in 130 accessions of *Lotus japonicus* and studied trait relationships, their genetic basis and identified two genes controlling accumulation of phosphate. Overall, our results exposed general patterns of phenotypic and metabolic responses to phosphate, as well as significant natural variation in these responses across Lotus accessions, which importantly were not necessarily related to the Lotus subpopulation classes (Fig. 1d,e). This indicates that population structure doesn’t confound to a large extent when studying responses to phosphate levels and don’t preclude screening for natural allelic variants that underlie these traits.

### Root responses to phosphate and the heritability of root and anion content traits

Low phosphate has been mostly associated with the inhibition of primary root growth and this process has been shown to be regulated by phosphate dependent iron toxicity in the Col-0 Arabidopsis reference accession. However, it is not frequently considered that other Arabidopsis accessions are not showing any inhibition of primary root growth upon phosphate starvation (Chevalier et al., 2003), a finding that had indicated that this response is not canonical. In line with that, phosphate deficiency dependent inhibition of early root growth was not observed in our Lotus panel as LP does only have a minor effect on Lotus primary root growth (Suppl fig. 2). This is consistent with previous reports for Lotus MG-20 ecotype (Volpe et al., 2016). Our data shows a broad variation of root growth responses among Lotus accessions, depending on phosphate availability. Altogether this points towards different adaptive strategies that have been selected in the Lotus natural populations in order to cope with phosphate starvation in their natural soil environments, in a similar manner to the natural variation of this response that has been described in Arabidopsis natural accessions.

Hierarchical clustering among accessions does depend on the phosphate level and not on the Lotus subpopulation (Fig. 1d-e), indicating that the observed responses to phosphate are not just an expression of the kinship of these accessions.

Beyond highly heritable traits, such as flowering time (Atwell et al., 2010) and seed dormancy (Kerdaffrec et al., 2016), whose variation strongly depends on plant adaptation to environmental conditions, in the last years different studies successfully used GWA for identifying genes and alleles regulating both plant nutrient concentration and root growth traits. By measuring plant cadmium (Chao et al., 2014), sulfur (Koprivova et al., 2013; Huang et al., 2016), sodium (Baxter et al., 2010) and phosphate (Kisko et al., 2018) tissue concentration, causal genes were identified. A similar approach has been used to trace and map root growth responses to iron deprivation (Satbhai et al., 2017), salt stress (Julkowska et al., 2017), zinc (Bouain et al., 2018), nitrogen (Gifford et al., 2013) and phosphate (Stetter et al., 2015) levels. In our attempts to integrate the two last approaches and recapitulate the natural variation of Lotus phosphate accumulation and root system architecture responses to phosphate levels, we observed a great variability among broad sense heritability between the two groups of traits (Fig. 2a). Interestingly, in our set up, the majority of traits related to anions showed higher BSH compared to root growth related traits, with the exception of primary root growth length. Lower BSH reflects a higher trait variance within genotype compared to the trait variance found across genotypes. We therefore expect that traits showing higher BSH are either highly responsive to the external environmental conditions and factors not taken into account in our experimental set-up, such as plate micropatterning or seed size, or that our measurement error was too high for those traits. Another possible reason for the difference of BSH between these trait classes could be the different number of replicates: for the RSA analysis, we analysed 8 biological replicates, whereas the anion content was based on 4 biological replicates. Nevertheless, the higher BSH did not result in larger number of significantly associated SNPs for single anion traits (Supplementary figure 4).

### Relation of phosphate content and root growth

During our investigation we found a surprising correlation between phosphate content and growth related traits exclusively in LP conditions (Fig. 2c): the longer the root, the less concentrated the phosphate. The most likely explanation for this seems to be that we observe phosphate dilution effect in which the limited amount of P that is available in the plant is distributed over a larger amount of tissue in case of larger accessions. Our initial observation on a broad panel of accessions, became even more evident when focusing on a single genetic background (Gifu), where less genetic confounding effect are present (Fig. 5). In this scenario, when plants are grown under low (20 μM) and mid (100 μM) phosphate, an even stronger and more significant negative correlation between plant biomass and plant phosphate concentration emerges. Again, this correlation is completely absent from plant grown under sufficient phosphate concentration (750 μM). It would not be surprising if a similar correlation could be observed for other main limiting factors for plant growth, such as nitrogen, sulfur or potassium. While it has been described in *Brassica oleracea* that shoot yield drives phosphorus use efficiency and correlates with root architecture traits (Hammond et al., 2009), this process seems not to have been described before. A possible reason this link was not previously described could be that the whole-single-plant resolution is usually missing. In fact, in experiments performed in Arabidopsis, many plants are usually bulked together before measuring any content and therefore extreme values are lost, therefore possibly occluding correlations. Conversely, in studies focusing on crop plants, due to the plant size, only a part of it is usually considered, therefore possibly missing organismal correlations. Given that nutrient levels are also responsible of a downstream cascade of gene transcription and cellular reprogramming (in the case of phosphate starvation they depend on PHR1 transcription factor), we think that plant biomass should always be taken into account when dealing with nutrient starvation condition, to avoid recursive confounding effects.

### Candidate genes for phosphate homeostasis

Phosphate is one of the main macronutrients and a limiting factor for plant growth, which is highly variable in natural and agricultural soils (Orgiazzi et al., 2018). We therefore expect a strong selection on plant genomes due to soil phosphate concentration and/or soil microenvironment (both biotic and abiotic). Nevertheless only few studies have identified causal genes involved in plant phosphate nutrition in the light of natural variation (for example Kisko et al., 2018; Stetter et al., 2015; Yang et al., 2012). By contrast, much more detailed knowledge has been acquired through forward genetics screening and transcriptomics approaches (mainly in Arabidopsis and rice) and subsequent validation of candidate genes.

Our GWAS analysis has detected hundreds of significant associations, among which are known regulators of plant root responses to low phosphate, such as *STOP1*. By combining candidate genes that were overlapping among traits, an approach that was similarly used in cereals (Chen et al., 2016), we selected and validated two of these, a Leucine-Rich-Repeat receptor kinase and a cytochrome B5 reductase. For each of these candidate genes, multiple LORE1 insertional mutants accumulate more soluble phosphate than the wt plants on high phosphate media (Fig. 4). However, despite the candidate genes being also associated with root traits, the mutants did not show aberrant root phenotypes. This could be due to various reasons: for instance, redundancy or genetic buffering might compensate for these genes for early root growth, the same loci might have a minor effect on RSA and stronger effect on phosphate levels and/or the same SNPs were in LD with other genes that could control RSA.

Despite the lack of early root phenotype, the clear involvement of these genes in the control of root phosphate concentration exposes two new phosphate regulating genes, which are among the first phosphate regulators known in Lotus. The Arabidopsis homologue of the cytochrome B5 reductase, CBR1 (Oh et al., 2016), was recently described as a crucial factor for iron uptake due to its role in activating plasma membrane H^+^-ATPase, responsible for acidification of the rhizosphere. CBR1 is involved in energy transfer at the ER level, it therefore could also control other important plant ion pumps that depends on electron potential. The inactivation of the Lotus homologue leads to the accumulation of phosphate in plant cells, even though the localization and the pool partitioning remains to be uncovered. *LAMP*, the LRR-RK is involved in regulating internal plant phosphate levels and might therefore, similarly to other plant membrane receptors that regulate nitrogen metabolism in Arabidopsis and/or rhizobial abundance in Lotus (Tabata et al., 2014; Okamoto et al., 2013), be involved in nutrient signalling.

We are aware that further functional studies are needed to mechanistically understand their role in phosphate uptake and/or recycling, eventually taking into account potential ligands such as regulatory peptides that are utilized for signalling during phosphate starvation.

## Material and methods

### Plant material and growth conditions

In total, 130 *Lotus japonicus* accessions were used (Shah et al., 2018). The names and accession numbers are listed in Supplementary Table 6. Seeds were scarified with sandpaper and then sterilized 14 minutes in 0.05% sodium hypochlorite. Subsequently, seeds were rinsed and washed 5 times in sterile distilled water. For the germination, seeds were positioned in imbibed filter paper, in sterile Petri dishes, and wrapped in aluminium foil. After 3 days at 21°C, young seedling were transferred to square plates (12 × 12 cm) containing growth medium (as described in (Giovannetti et al., 2017)). Both media used in this study were based on Long-Ashton solution (with two levels of phosphate concentration −20 or 750 μM, LP or HP, respectively-as in Supplementary Table 1) with 0.8% MES buffer (Duchefa Biochemie, Haarlem, The Netherlands), 0.8% agarose (to minimize phosphate contamination), and adjusted to pH 5.7 with 1M KOH. After adding the medium, plates were dried, closed, overnight in a sterile laminar flow hood. Two accessions, with four replicates per each accession, were placed on each plate. Each plate was replicated, with mirrored position of each accession to minimize any positional growth effects. Plates were placed vertically, and plants grown under long-day conditions (21°C, 16 h light/8 h dark cycle) with white light bulbs emitting 50 μmol/m^2^ /s. Every day the position of plates was shuffled to avoid positional effects.

### Analysis of root growth and anion content

Each day at the same time, for 9 days, plates were scanned with CCD flatbed scanners (EPSON Perfection V600 Photo, Seiko Epson, Nagano, Japan), and the images were used to quantify root parameters using Brat 2.0 (as described in Giovannetti et al., 2017). After 10 days, roots and shoots from 4 plants of each accession were weighed and frozen. For anion measurements in the initial screening, frozen plant material from 4 biological replicates was then homogenized in 1 mL of deionized water, and the anions- nitrate, phosphate and sulfate-were separated by the Dionex ICS-1100 chromatography system on an Dionex IonPac AS22 RFIC 4×250 mm analytic column (Thermo Scientific, Darmstadt, Germany) with 4.5 mM NaCO_3_/1.4 mM NaHCO_3_ as running buffer. LORE1 Lotus mutants were ordered from Lotus Base (Mun et al., 2016) and homozygous plants selected with specific primers (Supplementary table 9).

### Genome-wide association studies and overlap analysis

GWA mapping was conducted on the mean and median trait values using a mixed model algorithm (Kang et al., 2008), which has been shown to correct for population structure confounding (Seren et al., 2012), and using the homozygous SNP data from the Lotus accessions (Shah et al., 2018). SNPs with minor allele counts less than 10 were not taken into account. The significance of SNP associations was determined around the 5% FDR threshold computed by the Benjamini–Hochberg–Yekutieli method to correct for multiple testing (Benjamini and Yekutieli, 2001) and genes within a 10-kb genomic region spanning each SNP were considered, taking into account that LD decays to 0.2 in Lotus (Shah et al., 2018).

### Inorganic phosphate concentration measurements

Shoots and roots were collected, weighed and ground into powder in liquid nitrogen. The powder was incubated at 98°C in NanoPure water, for 1 hr, centrifuged for 20 minutes at maximum speed. Then 25 uL of a dilution 1:10 were used to determine inorganic phosphate concentrations using the molybdate phosphate assay (Sigma), following kit instructions, as previously described (Ames, 1966). Each 96-well plate contained a calibration curve for assessing phosphate concentration.

### Figures and statistical analysis

Data analysis and plots were conducted in Rstudio (RStudio Team, 2016) using the following packages: tidyverse, emmeans, UpSetR, corrplot, RColorBrewer, rmarkdown, multcompView and gplots. Plots were further modified for colours and layout in Adobe Illustrator CS6. All the scripts used to generate raw figures can be found (Supplementary file 1) and raw measurements data (Supplementary table 10). Number of replicates and statistical tests are indicated below every graph.

## Acknowledgements

The authors thank Samantha Krasnodebski for her help in amplifying Lotus accessions, Anna Małolepszy for the initial common effort on Lotus projects, Santosh Satbhai and the members of the Busch lab for the training in the Brat system and GWAS, Ümit Seren for assistance on Lotus GWAPP website, Patrick Hüther and Niklas Schandry for their help with R, Yasin Dagdas for further supervision and hosting in the lab, Irene Klinkhammer and Bastian Welter for helping with the anions’ quantification. Funding of this work was supported by the Austrian Academy of Science through the Gregor Mendel Institute, the Salk Institute for Biological Studies and a Marie Sklodowska-Curie Individual Fellowship, number 749044 (to MG). Research in SK’s lab is supported by the Deutsche Forschungsgemeinschaft (DFG, German Research Foundation) under Germany’s Excellence Strategy – EXC-Nummer 2048/1.

## Supporting data

Supplementary Figure 1. Sulfate and nitrate concentration is not dependent on plant size

Supplementary Figure 2. Lotus japonicus natural variation of root responses to phosphate media levels over time

Supplementary Figure 3. Lotus japonicus primary root length is negatively correlated with root phosphate concentration in plants grown under phosphate limitation

Supplementary Figure 4. Number of hits (p-value < FDR) per trait in anions vs. Brat analysis

Supplementary Figure 5. Cytochrome B5 reductase and LRR mutants accumulate more phosphate than wt in high phosphate media

Supplementary Figure 6. Plant phosphate concentration is negatively correlated with plant biomass when phosphate is a the limiting factor

Supplementary Figure 7. Cytochrome B5 reductase and LAMP mutants are not affected in root growth over different phosphate concentration

Supplementary Figure 8. Manhattan plots leading to the identification of cytochrome B5 reductase locus

Supplementary Figure 9. Manhattan plots leading to the identification of LAMP

Supplementary Figure 10. LIMIX model of GWAS leading to the identification of LAMP locus

Supplementary Figure 11. LIMIX model of GWAS leading to the identification of cytochrome B5 reductase

Supplementary Table 1. Modified Long-Ashton media solution

Supplementary Table 2. GWAS hit from LP:HP ratio of Lotus Brat results

Supplementary Table 3. GWAS hit from Lotus roots grown on LP media

Supplementary Table 4. GWAS hit from Lotus roots grown on HP media Supplementary Table 5. GWAS hit from Lotus anion content

Supplementary Table 6. List of Lotus japonicus accessions

Supplementary Table 7. Statistics summary for cytochrome B5 reductase and LAMP mutants

Supplementary Table 8. Top 500 SNPs from anion and root traits with p-value < 10E-5 Supplementary Table 9. Primers and mutant plants used in this study

Supplementary Table 10. Raw data from root system analysis and anion measurements

Supplementary Table 11. GO enrichment for GWAS hits of root system traits from LP media

Supplementary Table 12. GO enrichment for GWAS hits of root system traits from HP media

Supplementary Table 13. List of significant SNPs above benjamini-Hochberg FDR threshold

Supplementary file 1. List of R codes and plots used in this study (Rmd file).

